# The importance of Chargaff’s second parity rule for genomic signatures in metagenomics

**DOI:** 10.1101/146001

**Authors:** Fabio Gori, Dimitrios Mavroeidis, Mike SM Jetten, Elena Marchiori

## Abstract

An important problem in metagenomic data analysis is to identify the source organism, or at least taxon, of each sequence. Most methods tackle this problem in two steps by using an alignment-free approach: first the DNA sequences are represented as points of a real n-dimensional space via a mapping function then either clustering or classification algorithms are applied. Those mapping functions require to be genomic signatures: the dissimilarity between the mapped points must reflect the degree of phylogenetic similarity of the source species. Designing good signatures for metagenomics can be challenging due to the special characteristics of metagenomic sequences; most of the existing signatures were not designed accordingly and they were tested only on error-free sequences sampled from a few dozens of species.

In this work we analyze comparatively the goodness of existing and novel signatures based on tetranucleotide frequencies via statistical models and computational experiments; we also study how they are affected by the generalized Chargaff’s second parity rule (GCSPR), which states that in a given sequence longer than 50kbp, inverse oligonucleotides are approximately equally frequent. We analyze 38 million sequences of 150 bp-1,000 bp with 1% base-calling error, sampled from 1,284 microbes. Our models indicate that GCSPR reduces strand-dependence of signatures, that is, their values are less affected by the source strand; GCSPR is further exploited by some signatures to reduce the intra-species dispersion. Two novel signatures stand out both in the models and in the experiments: the combination signature and the operation signature. The former achieves strand-independence without grouping oligonucleotides; this could be valuable for alignment-free sequence comparison methods when distinguishing inverse oligonucleotides matters. Operation signature sums the frequencies of reverse, complement, and inverse tetranucleotides; having 72 features it reduces the computational intensity of the analysis.

## 1 Introduction

Metagenomics studies the genomic content of microbial communities, obtained through DNA sequencing technologies [1]. Essentially, a metagenomic dataset is a set of DNA sequences acquired from the genomes of an environmental sample. By bypassing the cultivation step, metagenomics is able to obtain microbial genomes unattainable through individual sequencing, since less than 1% of the microbes present in nature can be cultured [2]. Moreover, with metagenomics it is possible to infer the interactions occurring in a microbial community. Unfortunately, the potentialities given by metagenomics come with a price in terms of data analysis challenges: we do not know from which genome a sequence was sampled; in most of the cases, the full genomes of the community members are not available; even species number is unknown.

As a consequence, an important step of metagenomic data analysis is to detect to which organism, or at least to which taxon each sequence belongs to. This problem is tackled, for instance, by means of clustering methods (binning) [3], by prediction models constructed using available genomes (taxonomic assignment) [4,5], and by other similarity-based approaches that match sampled sequences with sequences in a database of reference [6,7].

Many of the binning and taxonomic assignment methods tackle the problem in two steps: first the DNA sequences are represented as points of a real n-dimensional space via a mapping function, then either clustering [3, 8–13] or classification algorithms [14, 15] are applied to these points. Typically, the mapping functions adopted in literature represent a given sequence with the frequency counts of a set of oligonucleotides: among these, tetranucleotides are the most used [8,9,12, 15]; sometimes frequencies of short oligonucleotides (length up to 6) are used together [11,14]; a few tools adopted oligonucleotides longer than 6 bases [3, 10, 16, 17]. More recent tools are based on tetranucleotides frequencies computed on contigs assembled from the reads of the metagenome [18,19].

In order to be effective for binning and taxonomic assignment, these mapping functions have to be *genomic signatures* [20]. A genomic signature is a mapping function that has the following properties: sequences sampled from the same genome are mapped to relatively similar points; sequences sampled from different genomes are mapped to significantly different points, and this difference is related to the phylogenetic distance of the source genomes. Signatures are also used for alignment-free sequence comparison [21]. The biological underlying explanation of this property of existing genomic signatures is still unclear. It is conjectured to be the result of more contributing factors [22], like GC content and phylogeny [23]; however the correlation between signature and phylogenetic distance appears to be not very strong, mainly due to the absence of divergence of oligonucleotide composition in some phylogenetically distant species [24].

Despite the relevance of genomic signatures for metagenomic data analysis, existing signatures were not designed to take into account the special properties of metagenomic data. Signatures are usually tested on sequences of 10,000 base pairs (bp) or more [20], while the sequencing technologies used for metagenomics generate sequences of 50–1,000 bp. Signatures for metagenomic data cannot be based on information extracted from source genome of a sequence, like many existing signatures do, since composition of the sequenced community is often unknown, and new species might also be present in the metagenome. Signatures need also to be effective with sequences containing errors, that can be generated by sequencing machines. Moreover, signatures for metagenomic data have to be effective even with sequences belonging to different strands of the genomes, because sequencing technologies might sample sequences from both strands.

Furthermore, the development of binning and taxonomic assignment methods was more focused on the algorithmic part rather than on the adopted mapping function/genomic signature. Attempts to introduce signatures for metagenomic applications are recent and validated the signatures on error-free sequences, sampled from a few dozens of species, and longer than real metagenomic sequences. For instance, the binning tool MetaCluster successfully tested and implemented a signature based on tetranucleotide frequencies ranking [13, 25], and showed its effectiveness for clustering metagenomic sequences. RAIphy used a signature computable on sets of sequences [16]; MetaProb computed normalized tetranucleotide frequencies on sequence sets [26]. Signature OFDEG was designed for error-free sequences of at least 8,000 bp [27]. Recently, ICOs signature was tested on 50,000 error-free sequences of 1,000 bp or more, sampled from 60 species [28].

In this study, we analyze and compare theoretically the statistical distributions of some existing and novel signatures; in particular, for each signature we study its dispersion among sequences of the same species and the dispersion of signature’s expected values among different species. A low within-species dispersion shows that the signature assumes similar values for sequences sampled from the same species. A signature’s expected value for a species gives us an indication of its average values for that species; high dispersion of species expected values indicates that the given signature maps sequences of different species to significantly different points.

Furthermore, we investigate the biological and statistical rationales that make signatures effective in dealing with metagenomics data. In particular, we take into account the effects of the generalized Chargaff’s second parity rule (GCSPR) on genomic signatures. This rule states that, in a given sequence of at least 50 kbp, an oligonucleotide and its reverse complement are approximately equally frequent [29,30]; the rule implies that the two oligonucleotides have approximately the same total count in a genome [31]. GCSPR rule applies to all the double stranded DNAs, organelles excepted [32].

We also provide a thorough comparative experimental analysis of those signatures on data more similar to the metagenomic ones than the ones used in signature studies [20, 27, 28], especially in terms of number of species, sequence length, and presence of errors. Our experimental setup is a significant extension of the one adopted in our previous work [33]: sequences have base-calling errors (1% rate); experiments are performed for multiple sequence lengths (150 bp, 500 bp, 1000 bp); we directly compare distributions of signatures’ distances instead of their mean values; the Area Under Precision-Recall curve (AUPR) replaces the Area Under ROC curve as an evaluation measure, to take into account that within-species sequence pairs are much fewer than between-species ones; the number of sequences pairs on which within-species signatures’ distances are computed has been increased hundred-fold.

The proposed signatures take into account the special properties of metagenomic data mentioned before. In particular, they are designed to be effective with sequences of the same genome sampled from different strands: these *strand-independent signatures* assume the same value for a sequence and its reverse complement, so that the source strand of the sequence become irrelevant for the signature. All the signatures we study are derived from the tetranucleotide frequencies signature; we will refer to this as the *standard signature*.

The most important novel signatures introduced in this study are the combination signature and the *operation signature*. Combination signature is a reordering of the features of the standard tetranucleotide frequencies signature; the adopted reordering makes combination signature strand-independent. Strand-independent signatures currently used in metagenomics [18,26] are derived form the *symmetrized signature*, that is obtained by summing the frequencies of reverse complementary tetranucleotides [24]. Therefore, combination signature proves that summing frequencies of inverse oligonucleotides, and thus loosing information, is not necessary to achieve a strand-independent signature.

The 72-feature Operation signature is instead obtained by summing the frequency of a tetranucleotide with the ones of its complement, reverse, and inverse. This was developed to study if exploiting GCSPR and the reverse, complementarity, and inverse relations between tetranucleotides [34] can reduce feature space dimension and increase error tolerance. In data analysis, in general, reduction of feature space dimension can be beneficial in many aspects: performances are less dependent from the data (see bias-variance trade-off [35]), the computational cost of the analysis is reduced, and the data can become more interpretable [36]. Dependency from data is a very important issue for metagenomic data analysis, because metagenomes are likely to contain novel microbes and communities: hence, an analysis method based on fewer features can be more accurate when facing these types of data. The reduction of computational cost is also relevant, because the size of metagenomes is growing so much that analysing them is already becoming expensive in terms of computational power and data storage [37].

We also define and test new signatures capturing the divergence from GCSPR. Since metagenomic sequences are much shorter than 50 kbp [29, 30] the hypotheses behind GCSPR are not completely satisfied and hence the frequencies of reverse complementary oligonucleotides may differ. These differences could change according to the taxonomic classification of the source genome, making them exploitable for our purpose: indeed, previous research showed that purine-pyrimidine asymmetry in mammalian mitochondrial DNA carries phylogenetic information [38].

## 2 Materials and Methods

### 2.1 Genomic Signatures and their Rationale

In this study we focus on signatures based on tetranucleotide (4-mer) frequencies, since previous works had demonstrated that these features carry a significant phylogenetic signal [39]. Let *w* denote one of the 256 tetranucleotides, represented as words of length 4 in the alphabet {A,C,G,T}; we denote by *w*^*RC*^ the reverse complement (also called *inverse)* of *w.* Note that 16 of these 256 tetranucleotides are *palindromic*, i.e, they coincide with their respective reverse complements *(w*^*RC*^ = *w*). A metagenomic sequence *s* is also represented as a word in the alphabet {A,C,G,T} but with no length limit; *s*^*RC*^ still denotes the inverse of *s.* For a given sequence *s*, we denote with *f_i_* and 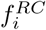 the frequencies with which tetranucleotides w_i_ and 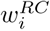 occur in *s*, respectively. We also denote with *w*^*C*^ and *w*^*R*^ the complement and reverse of *w*, respectively. Note that *w*^*R*^ and *w*^*C*^ are reverse complement of each other. Therefore, for the 8 distinct tetranucleotides coinciding with their reverse *(w* = *w*^*R*^), the relation *w*^*RC*^ = *w*^*C*^ holds; if *w* is palindromic, then *w*^*C*^ is another distinct palindromic (*w*^*C*^ = *w*^*R*^). The symbols 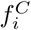 and 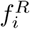 denote the frequencies of 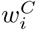 and 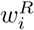 in a given sequence *s*, respectively.

We view a signature as a function *ρ*^*α*^ mapping a sequence *s* to a vector *ρ*^*α*^(*s*) = (*a1, …, a_n_*) of real numbers. A signature *ρ*^*α*^ is called *strand-independent* if it assumes the same value for each possible genomic sequence *s* and the sequence sampled from exactly the same genomic region but on the complementary strand, that is the inverse of s: that is, if the relation *ρ^α^(s)* = *ρ^α^*(*s*^*RC*^) holds for all possible sequences *s.*

First, we introduce the signature usually adopted in metagenomic data analysis:

*<u>Standard Frequencies Signature</u> ρ*^T^: This signature is defined by setting the *i*-th component *a_i_* of *ρ*^T^(*s*) to *f_i_*, for *i =* 1, …, 256. This signature has been used in many tools for metagenomic data analysis [8, 9,12,15]; among the analyzed signatures, it is the only one that is affected by the source strand of the sequence and does not exploit reverse complementarity of tetranucleotides. It is also affected by the deviation from GCSPR.

All the other signatures under examination in this work are strand-independent. The first group of signatures we examined are novel signatures that can be derived from *ρ*^T^ by simply reordering its features or selecting only some of them:

*<u>Minimal and Maximal complementarity signatures</u> ρ*^min^ *and ρ*^max^: These signatures are defined such that *ρ*^min^(*s*): = (*a*_1_, …, *a*_120_), with 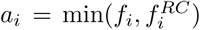 and 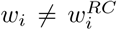, and *ρ*^max^(*s*): = (*a*_1_, …, *a*_120_), with 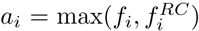 and 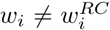. These signatures are affected by the deviation from GCSPR. Notice that in these signatures we employ only the 240 non-palindromic tetranucleotides.

*<u>Palindromic Signature</u> ρ*^P^: This signature considers only the frequencies of the 16 palindromic tetranucleotides (i.e.,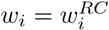). That is *ρ*^P^(s): = (*a*_1_, …, *a_16_)* with *a_i_ = f_i_.* The introduction of this signature is motivated by a study where the frequency distribution of palindromic tetranucleotides was shown to exhibit highest inter-species but low intra-species variance on 10,000 bp sequences [40].

*<u>Combination Signature</u>* (*ρ*^max^*, ρ*^min^*, ρ*^P^): As a combination of *ρ*^max^*, ρ*^min^ and *ρ*^P^, it maps a sequence *s* to a vector (*a_1_, …, a_120_, b_1_, … …, b_120_, c_1_, …, c_16_*), where the features 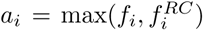 and 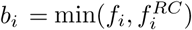 are derived from the non-palindromic 4-mers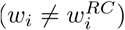, while c*_i_* = *f_i_* is computed for the 16 palindromic 4-mers. This signatures is actually a reordering of the 256 tetranucleotide frequencies composing *ρ*^T^, in a way that makes the signature strand-independent and is still affected by the deviation from GCSPR. As a matter of fact, this combination maps a given sequence *s* to a permutation of the 256 components of *ρ*^T^(*s*). Indeed, following the notation previously introduced, it can be proved that each feature of *ρ*^T^ corresponds to the frequency of a certain tetranucleotide *w.* If *w = w*^*RC*^ holds, than its frequency will be a feature of *ρ*^P^; if the equality does not hold, frequency of *w* will be a feature either of *ρ*^max^ or *ρ*^min^.

#### 2.1.1 Signatures exploiting generalized Chargaff’s second parity rule

Other signatures we examined reduce the number of features and increase error tolerance by exploiting GCSPR and other genomic symmetries; those signatures include a novel one called operation signature: *<u>Symmetrized Signature</u> ρ*^S^: This signature is obtained by summing the frequencies of distinct inverse 4-mers (see, e.g., [24],). It is defined as *ρ*^S^(*s*): = (*a*_1_, …*,a*_136_), with 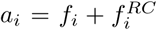 if 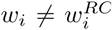, and *a_i_* = *f_i_*, otherwise. Notice that the vector *ρ*^S^*(s)* has 136 features, since 16 tetranucleotides are palindromic, i.e, they coincide with their inverse, and 240 are not (i.e., 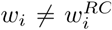. The symmetrized signature *ρ*^S^ can be seen as a simplification of the combination signature (*ρ*^max^, *ρ*^min^*, ρ*^P^), and hence *ρ*^T^, because 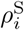 is equal to 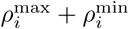 if 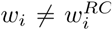 holds, otherwise 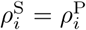. Indeed, a way to reduce the dimension of combination signature is to substitute some of its features with new features corresponding to their sum; the side effect of this approach is that it removes the distinction between the frequencies of the chosen tetranucleotides.

This signature exploits GCSPR to reduce feature space dimension. GCSPR states that, on sequences of 50 kbp or longer, *w_i_* and 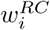 have approximately the same frequency, i.e. the relation 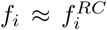 holds. Therefore, if sequences were sampled all from the same strand, we would just need to choose one of the two frequencies as features to build effective signatures. Since this is not possible, it is sensible to replace those two frequencies with their sum, that corresponds also to the sum of 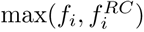 and 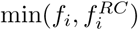. Indeed, thanks to the previous relation, we have that 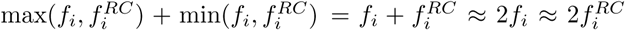 also hold; hence it is sensible to replace each of these 120 feature pairs with their sum 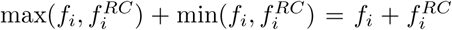, thus reducing the signature to 136 features. As a result, we obtain *ρ*^S^.

*<u>Symmetrized Rank Signature</u> ρ*^Rank^: This signature is defined such that *ρ*^Rank^(*s*) is the ranking induced by sorting the elements of *ρ*^S^(*s*) in descending order. This signature was used in recent works on metagenomic binning^1^ [13,25]; however, it was not specified how *ρ*^S^(*s*) ranking is performed when some ρ^S^(*s*) elements have the same value. We decided to perform a second ranking between features having the same values according to the alphabetical order of the respective tetranucleotides. For example, if the frequency of the palindromic 4-mer ’ACGT’ is equal to the sum of the frequencies of the reverse complementary pair ’AAAA’ and ’TTTT’, then the *ρ*^Rank^ value corresponding to the pair will be lower than the one of the single sequence, because ’AAAA’ precedes ’ACGT’ in the alphabetic order. Our choice of this second ranking was motivated by the simplicity of its implementation and computation.

*<u>Operation Signature</u> ρ*^O^: This signature is obtained by summing the frequency of a tetranucleotide with the ones of its complement, reverse, and inverse. It is inspired by a publication where the set of oligonucleotides is partitioned in equivalence classes with respect to complement and reverse operations [34]. It is defined as *ρ*^O^(*s*): = (*a*_1_, …, *a*_72_), with 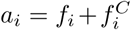 if 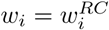 or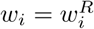, and 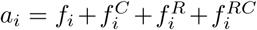 otherwise. Notice that the vector *ρ*^O^ (*s*) has 72 features: 8 features are given by the 8 sets 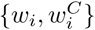 for which 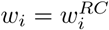 (and 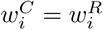); other 8 features are associated ot sets of the same form where (and 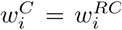); the remaining 224 tetranucleotides are all distinct, from their reverse, complement, and inverse, thus leading to 56 features.

As *ρ*^S^ is a reduction of *ρ*^T^, signature *ρ*^O^ can be seen as an additional simplification of *ρ*^S^, derived by exploiting complement and reverse relation between the tetranucleotides to further reduce feature space dimension. Indeed, *ρ*^S^ can be additionally reduced by substituting some of its features with new features corresponding to their sum, as we did before. Given a tetranucleotide *w*, we can observe that its complement *w*^*C*^ and the reverse *w*^*R*^ are one the reverse complement of the other. Therefore we can replace the 56 *ρ*^S^ feature pairs 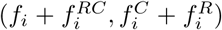 with their sum when the four tetranucleotides are distinct. 8 features 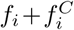 are obtained from the 16 palindromic tetranucleotides *w = w*^*RC*^ and *w*^*C*^ = *w*^*R*^ (e.g. *w* =’ACGT’, *w*^*C*^ =’TGCA’). The remaining 8 features of *ρ*^S^ corresponding to the *w* coinciding with their reverse (and hence *w*^*RC*^ = *w*^*C*^) are not changed (e.g. *w* =’ACCA’, *w*^*RC*^ =’TGGT’).

#### 2.1.2 Signatures capturing deviation from generalized Chargaff’s second parity rule

The following novel signatures were designed to capture solely the deviation from GCSPR in the given sequence. The deviation is computed with respect to tetranucleotide frequencies; their features are derived from the non-palindromic tetranucleotides:

*<u>Asymmetry Signature</u> ρ*^A^ : This signature is defined as *ρ*^A^ (*s*): = (*a*_1_, …, *a_120_*), where 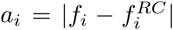. This is the only signature that measures the deviation with respect to sequence length, because of the way *f_i_* is defined.

*<u>Skew Signature</u> ρ*^Skew^: This signature is based on the standard relative skew index usually adopted in literature, such as [41]. It is defined as *ρ*^Skew^(s): = (*a*_1_, …, *a*_120_), where

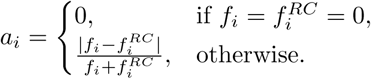

*<u>Ratio Signatures</u> ρ*^Ratio1^ *and ρ*^Ratio2^: These signatures are defined for the 4-mers that have different reverse complement as *ρ*^Ratio1^(s) = (*a*_1_, …, *a*_120_) and *ρ*^Ratio2^(s) = (*b*_1_, …, *b*_120_), where

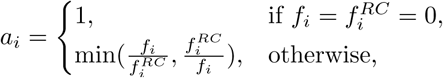

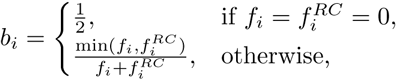

for 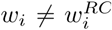.

*<u>JS Signature</u> ρ*^JS^: This signature is based on Jensen-Shannon divergence [42], and is defined as *ρ*^JS^(*s*) = (*a_1_*, …, *a*_120_) with

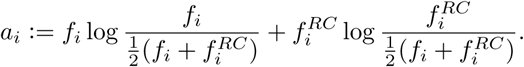

This signature is based only on the non-palindromic tetranucleotides.

Similarity between signatures was computed by using *L_1_* distance (also known as Manhattan distance). The choice of this distance is motivated by its use in previous methods for taxonomic assignment of metagenomic sequences [12,19]. Moreover, the distances most often used in literature on genomic signatures are based on *L_1_* multiplied by an averaging factor [20,24,43]. Given a genomic signature *ρ*^*a*^, the related signature distance between two nucleotide sequences s, *z* is defined by computing the *L_1_* distance between *ρ^a^(s)* and *ρ^a^(z).*

We also analyzed a few combinations of pairs of signatures, such as *(ρ*^S^ (*s*), *ρ*^A^ (*s*)), *(ρ*^min^*(s), ρ*^max^(*s*)), and the remaining combinations of *ρ*^min^*, ρ*^max^ and *ρ*^P^. Similarly, we studied the combinations 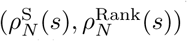 and 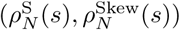, where 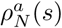 is the normalized version of a given signature *ρ^a^(s);* 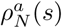 is defined as *ρ^a^(s)* divided by the maximum value that can be achieved by the related signature distance. This maximum distance depends on the sequence length. The maximum values of the signatures are provided in Supplementary Material (Table 1). Performances of the sum of minimal and maximal complementary signatures, namely *ρ*^min^ + *ρ*^max^, were also analyzed.

### 2.2 Data acquisition and preprocessing

Complete genomes of 1,284 prokaryotes were downloaded from the NCBI ftp server^2^. The list of the genomes is provided online^3^.

From a given genome, three sets of sequences were randomly sampled from both strands, simulating a sequencing error of 1%. Each of these sets consists of 10,000 possibly overlapping sequences with same length. Three sequence lengths where considered: 150 bp, 500 bp, and 1,000 bp. A sequence of length *l* is sampled by copying a random sub-sequence of *l* consecutive bases from a strand of a genomic sequence; only sequences made exclusively of the four bases {A,C,G,T} were considered. Sequences were sampled with a random base-calling error of 1%. The sequencing error was simulated with the method adopted in [13]: each base had 1% of probability of been wrongly sequenced. The probability was uniformly distributed among the other three bases (e.g A has 99% probability of being correctly sequenced as A; A has a probability of 0.33% of being sequenced as C,G, or T, separately). This simple error model was chosen to make results not affected by biases of specific sequencing technologies. The NCBI taxonomy^4^ [44] was used as reference taxonomy of the analyzed prokaryotes. Sequence sampling, analysis of the results and plotting were carried out using the following Python packages: Biopython [45], SciPy [46], IPython [47], and Matplotlib [48].

### 2.3 Computing signatures values

Revising the methodology employed in related works [13], we generated sets of sequences and evaluated the dissimilarity of the signature values on pairs of these sequences. Specifically, we evaluated the quality of a signature based on its property of assuming similar values for sequences of the same genome, and different values for sequences of different ones. We also evaluated the signatures’ performance at taxonomic levels, by considering the *taxonomic distance* of two sequences as the taxonomic rank of the lowest common ancestor of their source genomes in the taxonomy tree.

Specifically, for each of the three sequence lengths (150 bp, 500 bp, 1,000 bp) we created 9 sets of sequence pairs, where each set corresponds to a different degree of diversity of the source genomes. Subsequently, signature distances between sequences for each pair of the sets were computed. From the resulting distance values, 9 distributions of distances were obtained for each signature. A first distribution was generated using the distances between sequences of a same genome *(intra-genome* signature distances): for each genome, we computed all the pairwise signature distances between the 10,000 sequences of that genome. These ∼ 6.42· 10^10^ distances (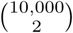 sequence pairs for 1,284 genomes) provided a distribution of intra-genome distances for the given signature. Each distribution was stored as a histogram of distance frequencies.

The other 8 distributions of distances were generated by computing distances between sequences from different genomes *(inter-genomic* signature distances), where each of the 8 distributions was obtained by considering a different level of taxonomic distance of the compared genomes. Specifically, we created 7 sets of organism pairs, one for each of the following taxonomic ranks: Species, Genus, Family, Order, Class, Phylum, Superkingdom. The set of pairs associated to a rank r consisted of 1,000 different pairs of organisms randomly selected among those whose lowest common ancestor in the taxonomy tree was at rank r. For each pair of these organisms, we randomly selected 1,000,000 pairs of genomic sequences from the set of all the sequences sampled from these genomes, and calculated the resulting distances. These 10^9^ distances (1,000,000 sequence pairs for 1,000 genome pairs) provided a distribution of inter-genomic distances at rank r. Each distribution was stored as a histogram of distance frequencies, whose bins were the same used for intra-genome distances histogram. Furthermore, we also created a set of organism pairs where each element is made by a *bacterium* and an *archaeon*, the two superkingdoms of the *Prokaryotes.* We computed and stored a signature distance distribution for this set of organism pairs using the same methodology applied for the other inter-genomic distances. We refer to this distribution as the inter-genomic signature distance distribution at prokaryotes level.

### 2.4 Evaluating the effectiveness of signatures experimentally

We assessed the capability of a genomic signature to preserve the taxonomic relations between the source genomes of pairs of sequences. Specifically, for each genomic signature, we tested if the related signature distance yielded small values for sequence pairs of taxonomically closely related source microbes, and greater values for sequences of distantly related microbes. To this aim, the signature distance was considered as a score for the sequence pair; the score quantifies the degree of relation between the source genomes, according to the given signature. Higher scores correspond to sequences that are more likely to belong to taxonomically distant genomes, according to the related signature.

Performances of different signatures were compared through Precision Recall (PR) curves [49]. The performance of each signature was evaluated at different *representation levels:* Intra-genome, Species, …, Superkingdom. For a given representation level, sequence pairs were partitioned in two sets: the pairs having taxonomic distances up to the associated level, called “positives”, and the remaining pairs, the “negatives”. Specifically, for intra-genome representation level, the set of positive sequence pairs was made by the pairs sampled from the same genome; the remaining pairs formed the set of negatives. For representation level corresponding to taxonomic rank *r*, instead, we considered as positives all the sequence pairs such that the lowest common ancestor of the taxa of their source genomes was at rank *r* or lower. The remaining pairs were the negatives. Having defined the set of positives and negatives, we could compute the set of “true positives” and “false positives” for a given signature distance threshold and derive the PR curve. Given a signature distance threshold, we considered as “true positives” the positive pairs whose distance was below or equal to the threshold; similarly, “false positives” were made by the negatives with distance below or equal to the threshold. Therefore, for each distance threshold *t* we could compute the Precision and Recall, defined as follows:

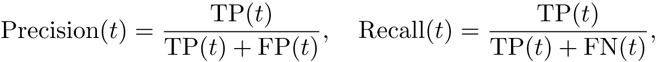

where FN(t), TP(*t*), and FP(*t*) indicate the number of false negatives, true positives and false positives for *t*, respectively. Plotting Recall on the x-axis and Precision on the y-axis, a point in the PR space is derived for a given *t.* Varying *t* among the values of our distance distributions, we produced the PR curve for the associated signature. We used the PR curve because it can clearly show if a signature *ρ*^*α*^ is always better than signature *ρ*^*β*^, namely if the PR curve of *ρ*^*α*^ is always above the one of *ρ^β^.* PR curve was preferred to Receiver Operating Characteristic curve because it is more informative when data are highly skewed with respect to negatives/positives abundances [49]. This property is relevant in our case, since in a generic metagenomic dataset the sets of sequence pairs corresponding to different levels of taxonomic diversity can have different sizes; for instance, it might happen that pairs belonging to related genomes, i.e. the positives, would be much fewer than pairs of distantly related ones. As an index of signature quality, we used the Area Under the PR Curve (AUPR) [49].

PR curves were derived from the histograms of distance frequencies previously obtained, simulating the signature performances on three different community structures. The community structures are given by the topology of the taxonomic tree of the community members. The first community structure, called *complex*, is a binary taxonomic tree: the taxa of each rank have two descent taxa, and each species has two distinct strains in the community. As shown in Supplementary Material (Section 2.2), with this structure the number of sequence pairs with taxonomic distance at a given rank increases exponentially with rank highness. We decided to study such a complex structure because it is known that binning methods have problems with communities made by many species. The second community structure, called *medium*, is made by a total of 11 strains distributed among 7 species (Supplementary Figure 1); 6 of these species belong to the same phylum. In this structure, there are no sequence pairs with taxonomic distance at rank Class and Superkingdom, because no species pair has lowest common ancestor at these ranks. The third community structure, called *simple*, is made by a species with 3 strains and by other 3 species with one strain each (Supplementary Figure 2). These 4 species belong to 3 phyla of the same superkingdom. In this structure, the sequence pairs are present only for taxonomic distance at intra-genome level and at ranks Class and Superkindom. The detailed description of these structures is available in the Supplementary Material. To analyze the effectiveness of a signature on a given community structure, the histograms of distance frequencies for the different ranks were rescaled: this was done to take into account that, for a given structure, the numbers of sequence pairs corresponding to different levels of taxonomic diversity of the source genomes respect a certain distribution. Details about the rescaling are provided in the Supplementary Material (Section 2.1).

For a given representation level and community structure, two histograms of distance frequencies were derived from the rescaled histograms; these two histograms represented the distances for the positive and the negative sequence pairs, respectively. For intra-genome representation level, the rescaled histogram related to intra-genome distances gave us the distance frequencies for the positive sequence pairs, the ones sampled from the same genome. The remaining rescaled histograms were added bin by bin, giving us the distance frequencies for the negative sequence pairs. For representation level corresponding to rank *r*, the rescaled histograms related to taxonomic distance at rank *r* or lower were added bin by bin, giving us the distance frequencies for the positive sequence pairs, i.e., the ones such that the lowest common ancestor of the taxa of their source genomes was at rank *r* or lower. The distance frequencies for the negatives were obtained in an analogous way, using the remaining rescaled histograms.

For each signature, the PR curve was derived from the histograms of positive and negative signature distances, respectively; the histograms shared the same bins. The PR curve was produced varying the threshold *t* among the edges of the histograms. Given a histogram, the number of sequence pairs whose distances were lower or equal than *t* was computed adding the histogram values for bins whose edges were lower or equal than *t.* Similarly, the number of sequence pairs whose distances were higher than *t* was computed using the histogram bins higher than *t.* Therefore, from the histograms we could compute the number of positives, true positives and false positives and hence the PR curve.

## 3 Results and Discussion

### 3.1 Theoretical analysis on the effect of GCSPR on signatures

We analyze and compare theoretically the statistical distributions of seven signatures; in particular, we study the signature dispersion among sequences of the same species (that should be as low as possible) and the dispersion of its expected values among different species (that should be as high as possible). The seven signatures under examination in this section are: four strand-independent signatures, namely the standard signature *ρ*^T^, maximal *ρ*^max^ and minimal *ρ*^min^ complementary signatures, combination signature (*ρ*^max^, *ρ*^min^, *ρ*^P^); two GCSPR-based signatures, namely symmetrized signature *ρ*^S^ and operation signature *ρ*^O^; and asymmetry signature *ρ*^A^, capturing the deviation from GCSPR.

We model signatures’ features of sequences as random variables. By analyzing the statistical dispersion of random variables corresponding to signature feature, we can assess theoretically the intra-species dispersion of signatures: a lower dispersion would correspond to better results, because it means that the signature assumes similar values for sequences sampled from the same species. We also analyze the inter-species discrimination capacity of signatures by looking at their distributions of per-species expected values: in this case, a higher dispersion indicates a better performance of the signature, because overall the signature values for different species will be more distant between each other.

Let *V_i_* be the random variable corresponding to the total occurrence of k-mer *w_i_* in a sequence randomly sampled from a given strand of a given organism, for *i* = 1, …,4*^k^*. Consistently with literature [26], we assume that *V1, …, Vi, …, V_4_^k^* follow a multinomial distribution with success probabilities *g_1_, …, g_i_, …, g_4_^k^*, respectively. It is sensible to take *g_i_* as the total occurrence of *w_i_* in the given strand divided by the genome size of the given organism; in particular, the sum of the success probabilities must be equal to one:

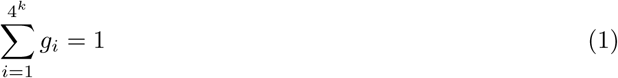

In particular, each *V_i_* follows the binomial distribution B(*g_i_,n*), where *n*: = *l* – *k* + 1 is the number of subsequences of length k in a sequence of length *l*. For small values of *k* (e.g. *k =* 4) it is sensible to assume that all the *g_i_* are strictly positive *(g_i_* > 0) because each k-mer will occur multiple times in the genome.

We now focus our attention on the random variable *X_i_*: = *V_i_/n*, that is the relative frequency of k-mer *w_i_* in sequences sampled from a given strand of a given organism. It is plain to see that *X_i_* corresponds to the *i*-th feature of standard signature 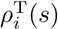. By analyzing the statistical dispersion of *X_i_*, we can assess whether the feature of *ρ*^T^ corresponding to *w_i_* tends to assume the same values for all the sequences sampled from the same strand of the same organism. We denote by 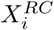 the variable corresponding to the relative frequency of the reverse complement 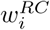.

GCSPR indirectly explains why standard signature *ρ*^T^ is effective on metagenomics data despite being strand-dependent; it also allows us to study *ρ*^T^ for sequences sampled from different strands. As recently stressed, GCSPR implies that inverse k-mers have approximately the same total count in a genome [31] and therefore we can assume that the global frequencies of *w_i_* and 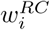 are equal:

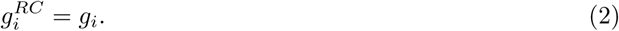

Let *s* and *z* be two sequences sampled from the opposite strands of the same genome. To determine the similarity between the two sequences, 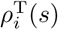 actually should not be compared to 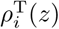 because the latter contains the relative frequency in *z* of the reverse complement of *w_i_* in the strand of *s*, namely 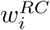. However, equation (2) implies that *X_i_* and 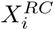 are identically distributed, and hence the relative frequencies of *w_i_* and 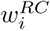 in sequences of the same genome follow the same probability distribution, irrespectively of the source strand (see Supplementary Material Section 1). Therefore it is sensible to compare 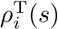 and 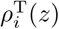; in particular the two paired features have identical mean and variance:

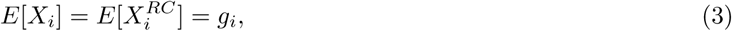

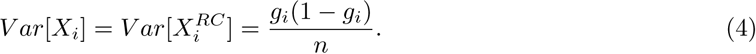

By assuming that word frequency *g_i_* is approximately the same in all the organisms of the same species, we can extend our theoretical analysis of *ρ*^T^ to all the sequences sampled from the same species. For standard signature, as for most of the signatures we analyze, an increase in sequence length *l* leads to lower variance; this is sensible for *X_i_*, because the longer the sequence, the narrower will be the difference between local and global frequencies of k-mers. For the rest of the manuscript, we assume that the indices of non-palindromic *w_i_* are ordered such that 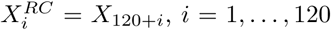; the last 16 random variables *X*_241_, …, *X*_256_ correspond to palindromic k-mers. Thanks to this ordering and to equation (2), the constraint (1) can be rewritten as:

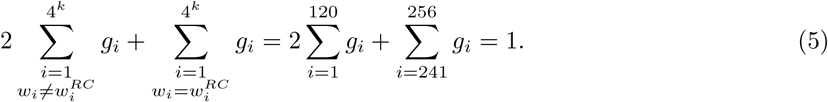

The expected value of random variable *X_i_* associated to non-palindromic k-mers therefore cannot be higher than 1/2.

As measures of intra-species dispersion of the *i*-th feature, we use the coefficient of variation (CV), defined as the ratio between the standard deviation and the mean of that feature for sequences sampled from the same species. For standard signature *ρ*^T^ the coefficient of variation is as follows:

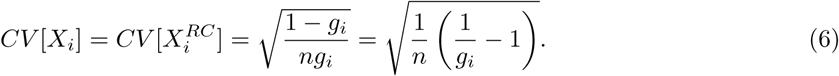

Signature with features having lower intra-species coefficient of variations have lower dispersion, and therefore are better because they tend to assume similar values for sequences of the same species. To compare the inter-species discrimination power of signatures, first of all we rescale the distribution of the expected signature value to the same range [0,1]: each feature is rescaled such that *ρ_i_*(0) = 0 and *ρ_i_*(1/2) = 1. If two signatures have the same distribution of species expected values for the *i*-th feature, then they have the same inter-species distinction power. Otherwise we look at the coefficient of variations of the random variable corresponding to the expected value of the *i*-th feature for a given set of organisms; in this case, the higher the coefficient of variation, the higher is the discrimination power.

#### 3.1.1 Symmetrized signature has lower intra-species dispersion than standard signature

Symmetrized *ρ*^S^ and operation signature *ρ*^O^, as all the signatures based on the sum of features of the standard signature *ρ*^T^, have smaller intra-organism coefficient of variation than *ρ*^T^ (Supplementary Information Sections 1.1 and 1.4). That result is obtained by modelling the sum of features as a random variable Ù*_j_* in the following way:

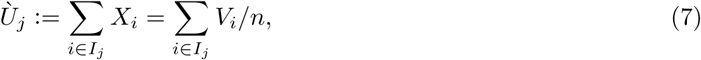

where *I*_1_, …,*I_h_* are a partition of the index set {1, …,4*^k^*}; for example, for *ρ*^S^ and *ρ*^O^ the set *I_j_* corresponds to the indices of words 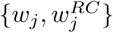 and 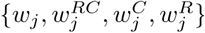, respectively. It is worth to mention that the use of GCSPR was not required for this observation.

Nevertheless, summing features could reduce the inter-species discrimination efficacy of a signature: sequences *s_1_* and *s*_2_ sampled from different organisms could have similar values for *Ù_j_* despite having distinct *g_i_*’*s.* For example, let us suppose that a signature has a feature given by the sum of the first two features of *ρ*^T^, and that *X*(*s_1_*) = (1/128,1/64, …) and *X*(*s_2_*) = (5/256,1/256, …). We have that the inequality (*X*_1_(*s*_1_), *X*_2_(*s*_1_)) *≠* (*X*_1_(*s*_2_), *X*_2_(*s_2_*)) holds but *X_1_*(*s_1_*) + *X*_2_(*s*_1_) = *X_1_(*s*_2_)* + *X*_2_(*s*_2_).

However, GCSPR gives symmetrized signature *ρ*^S^ the same inter-species discrimination power of the standard signature *ρ^T^.* Our analysis therefore indicates that signature outperforms standard signature because it has a lower intra-species dispersion but the same inter-species discrimination power. To illustrate that, by following equation (7) we model 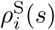 as a random variable as follows:

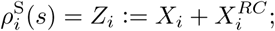

as shown at the beginning of Section 3.1, GCSPR implies that *X_i_* and 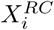 are identically distributed and have mean and variance as in equations (3) and (4). Therefore *Zi* has the following mean, variance, and coefficient of variation (see Supplementary Material Section 1.1):

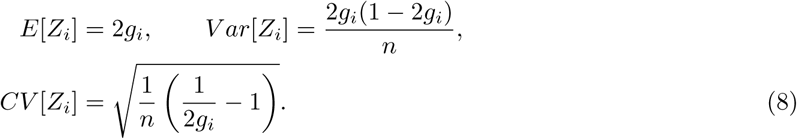

We compare the inter-species discrimination capacities of symmetrized and standard signatures by looking at their distributions of per-species expected values. For the 240 non-palindromic k-mers, symmetrized signature has 120 features because it combines the values of *X_i_* and 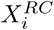. Therefore, we compare the distributions of the expected values of *Z_i_* with the distribution of the average expected values of *X_i_* and 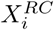. By the definition of *Z_i_*, we have that the expectation of *Z_i_* is twice the average expectation of *X_i_* and 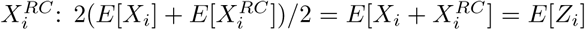. Since *Z_i_* takes values between zero and one, 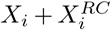 is the rescaling of 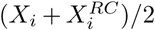 on that range and has the same expectation of *Z_i_.* Since the expected values are identical for the same species, the distributions of their expected values per-species are also identical.

Inter-species distinction reduction could actually occur for operation signature, although it is partially compensated by GCSPR. Let 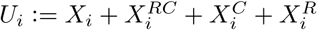 be the random variable corresponding to 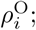; as observed at the beginning of Section 2.1, 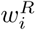 is the inverse of 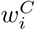 and therefore the GCSPR implies that *U_i_* follows a distribution with mean 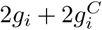 and variance 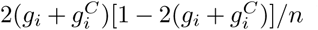 (Supplementary Material Sections 1.1). This means that the random variable corresponding to the sum of four k-mers actually depends on only two of them; however, it is likely that summing the frequencies of *w_i_* and 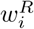 will have negative effects on the performance of the signature, because there is no symmetry rule for those kind of pairs. Specifically, operation signature cannot distinguish organisms having different values for k-mer frequencies of reverse words *g_i_* and 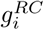 but the same value for the sum 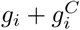.

#### 3.1.2 Combination signature performs in between standard signature and symmetrized signature

To simplify the analysis of combination signature, in this section we consider the normal approximation of *X_i_*. Specifically, we replace each binomial variable *V_i_* with its normal approximation 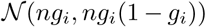. Consequently, even the variable *X_i_* follows a normal distribution because it is now the product of a constant term 1/*n* and a Gaussian variable; its mean is *g_i_* and its variance is *g_i_*(*1 – g_i_*)*/n*:

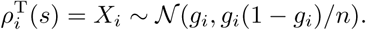

Let 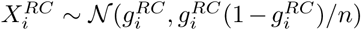 be the frequency of 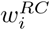 in sequence *s.* Given the normal distribution of *ρ*^T^ features, it follows that maximal 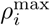 and minimal 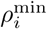 features follow the distribution of the maximum and the minimum of the pair of Gaussian variables 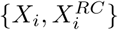, respectively. We can therefore define the following random variables corresponding to the signature values:

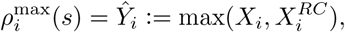

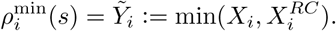

Formulas for first and second moment of maximum and minimum of pairs of Gaussian variable are known [50] and from those we can express mean and variance of 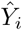 and 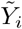 (see Supplementary Material 1.2). Since those maximal and minimal complementarity signatures are strand-independent, random variables 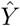 and 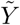 and their moments are also strand-independent.

The GCSPR allows us to show that maximal and minimal signature features, and thus the features of combination signature, have lower variance than the standard signatures. Equation (2) indeed drastically simplifies the first and second moment of 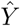 and 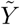 and hence *ρ*^max^ and *ρ*^min^. By replacing 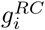 with *g_i_* we obtain (see Supplementary Material 1.2):

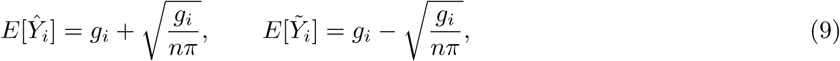

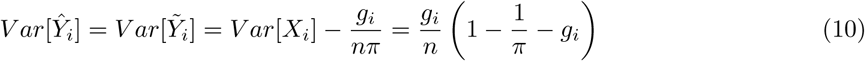

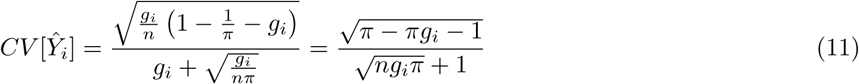

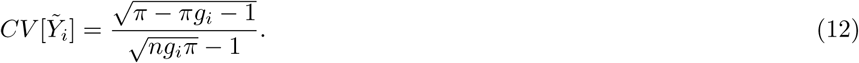

Since *g_i_/nπ* is strictly positive, it is plain to see from (10) that maximal and minimal signature features have lower variance than the standard signature features for all the non-palindromic k-mers – for the palindromic there is no difference.

The combination signature has lower intra-species dispersion than the standard signature but higher than the symmetryzed one (Figure 1). For the 16 palindromic k-mers, standard, symmetrized, and combination signatures are identical. Since combination signature for the 240 non-palindromic k-mers is a combination of maximal and minimal complementarity signature, the coefficients of variation of its features do not have the same formula. Therefore, we study the average value of 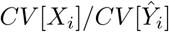 and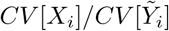. As shown in Section 1.4 of Supplementary Material, this is higher than one, and therefore combination signature has a lower dispersion than standard signature. On the other hand, in the same section we show that combination signature has higher dispersion than symmetrized signature for the non-palindromic k-mers, as shown in Figure 1:

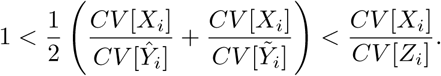

Combination signature has the same inter-species discrimination power of the standard signature. For the 240 non-palindromic k-mers, combination signature has 120 pairs of features, namely 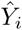 and 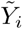, and each of these pairs is a function of the values of *X_i_* and 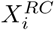. Therefore, we compare the distributions of the average expected values of 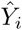 and 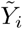 with the distribution of the average expected values of *X_i_* and 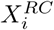. Thanks to equations (9) we have the following equality: 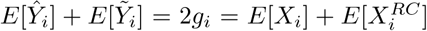. Hence the distributions of 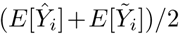 and 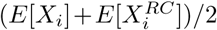 are identical, and the two signatures have the same inter-species discrimination power. Since the expected values are identical, the distributions of their expected values are also identical.

**Figure 1.**
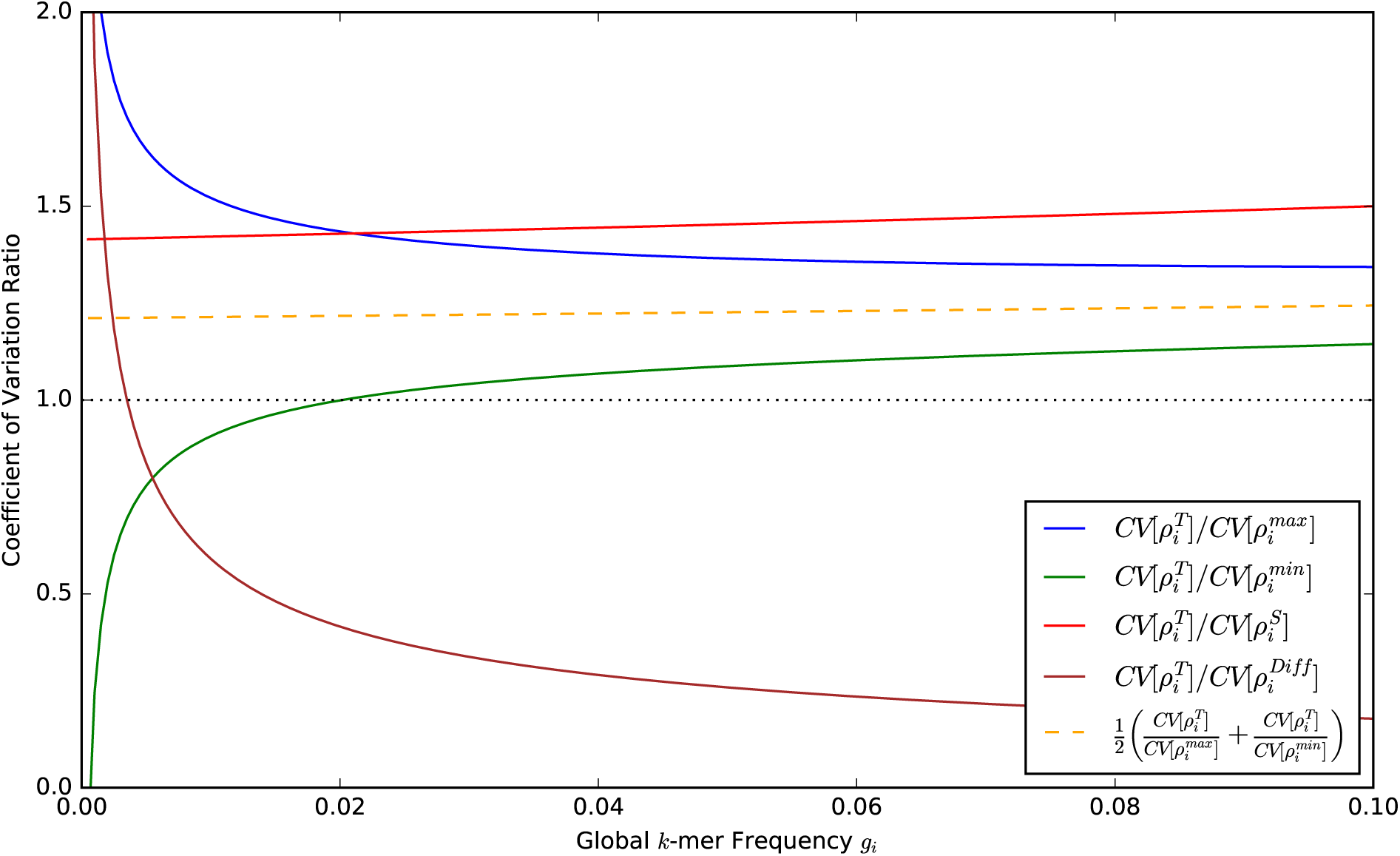
Ratio between coefficient of variations of some studied signatures and the coefficient of variation of the standard signature *ρ*^T^ for sequences of 500bp. Relevant ratio are displayed for symmetrized signature, combination signature, and asymmetry signature

#### 3.1.3 Divergence from GCSPR carries a phylogenetic signal

The expected values of asymmetry signature *ρ*^A^ are functions of the global word frequencies, and therefore *ρ*^A^ carries a phylogenetic signal. Indeed the *i*-feature of asymmetry signature is defined as the absolute difference between the frequencies of *w_i_* and 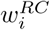; we show that the random variable *A_i_* corresponding to the value of 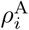 among sequences sampled from the given organism can be rewritten as a function of 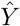 and 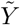:

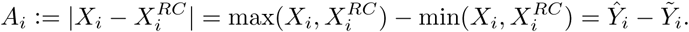

The mean, the variance, and the coefficient of variation for this signatures are (see Supplementary Material section 1.3):

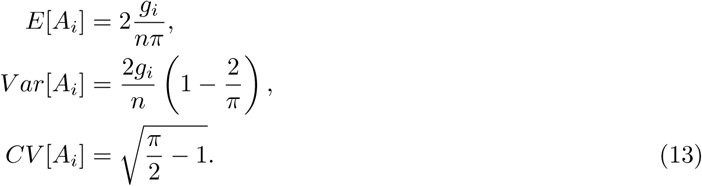

in contrast with all the other signatures studied, its intra-species dispersion does not decrease with read length and is actually constant (13).

The intra-species dispersion seems mostly worse than the symmetrized signature, and probably also of the plain signature. Indeed, the intra-species coefficient of variation strictly decreases with k-mer frequencies; it was lower than standard and symmetrized signatures for frequencies above 0.00351 and 0.00175, respectively (Figure 1).

We conduct an analysis of the intra-species coefficient of variation. Let *F_i_* be the random variable corresponding to the expected values of k-mer *w_i_* frequency for an organism belonging to a given set of organisms. The coefficient of dispersion of *F_i_* measures the intra-species dispersion of the standard signature: the higher this coefficient, the more spread are the expected value for feature *i* of the standard signature. Therefore, a higher coefficient leads to better discrimination between the given organisms. Our analysis shows that asymmetry signature can outperform the standard signature intra-species discrimination power if and only if 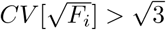 (see Supplementary Section 1.5).

#### 3.1.4 GCSPR-based signatures have better error-tolerance

An advantage of symmetrized signature over combination signature is in its tolerance to local deviation from GCSPR. In particular, symmetric deviations from k-mers frequence parity will have no effect on symmetrized signature. Indeed, it is known that Chargaff’s second parity rule, stating that complementary nucleotides have the same frequency along a strand, may not hold for sequences shorter than a species-dependent “critical fragment length”; this length is comprised between 6 kbp and 50 kbp [51]. The parity rule seems the result of alternating regions with different signs of deviation from parity. Consequently, it is reasonable to assume that even the version of the rule generalized for k-mers (the GCSPR) will not hold for some of the reads, whose length is for sure lower than the GCSPR-equivalent of the critical fragment length. For sequences of those regions, *X_i_* will still follow a binomial distribution but with a success rate *ǵ_i_* = *g_i_* + *δ_i_*, with *δ_i_* ∈ (– *g_i_*, 1/2 – *g_i_*) dependent on the sampled region and such that 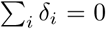. Let us focus on a special case of this deviation from parity here called *symmetric deviation*, where complementary k-mers frequency deviate from parity but with opposite deviation of the same size, i.e. 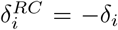. Symmetrized and operation signature will be uneffected by this deviation because the sum of the success rates of complementary k-mers will be the same: 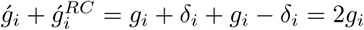

By summing frequencies of different tetranucleotides, the GCSPR-based signatures are more tolerant than the standard signature with respect to base-calling errors. It is plain to see that symmetrized signature *ρ*^S^ has some degree of tolerance to base-calling errors: if base-calling errors replace a tetranucleotide *w* with its inverse *w*^*RC*^, then the sum of their frequencies will be preserved. Moreover, if a base-calling error replace tetranucleotide *w_i_* with 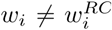, then the features of *ρ*^S^ corresponding to them could still be preserved not only if the opposite error occurs (*w_j_* replaced by *w_i_)* but also if any of *w_j_* and 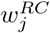 is replaced either by *w_i_* or 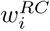. Those two mechanisms of error-tolerance of *ρ*^S^ are strengthened in *ρ*^O^. Indeed, feature associated with tetranucleotides *w, w*^*C*^, *w*^*R*^, *w*^*RC*^ will be preserved not only if a base-calling error replace one of those with its inverse, but also if any of the them is replaced by one of the other three. Moreover, if a base-calling error replace tetranucleotide *w_i_* with 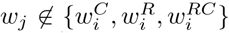, then the features of *ρ*^O^ corresponding to them could still be preserved if any of 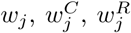, and 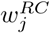 is replaced by any of 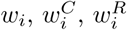, or 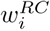 (hence there are 16 possible compensating errors instead of 4 and 1 of *ρ*^S^ and *ρ*^T^, respectively).

### 3.2 Experimental results

Signatures’ experimental performances were assessed through Precision Recall curves, using the Area Under the Precision Recall curves (AUPRs). The AUPRs obtained by a signature for different levels of representation were compared with the ones of an artificial signature that cannot distinguish between the different taxonomic ranks, because it has the same distance distribution for each rank. We show in the Supplementary Material that the AUPR of this artificial signature is equal to the ratio of positives in the data. If the AUPR of a signature is higher than the one of this artificial signature, then it can be considerate efficacious; the higher the AUPR, the better the signature.

Experimental performances of symmetrized, combination, and standard signatures were consistent with the theoretical analysis. Indeed, symmetrized signature *ρ*^S^ outperformed combination signature (*ρ*^max^,*ρ*^min^,*ρ*^P^), which in its turn outperformed standard signature *ρ*^T^; these relations held for each representation level and community structure. More in general, the best experimental results were achieved by strand-independent signatures summing or reordering most of the tetranucleotides frequencies, like *ρ*^S^, (*ρ*^max^*,ρ*^min^,ρ^P^)*, ρ*^min^ *+ ρ*^max^, and (*ρ*^max^*,ρ*^min^) (Figures 2, 3, 4, and Supplementary Figure 4).

**Figure 2.**
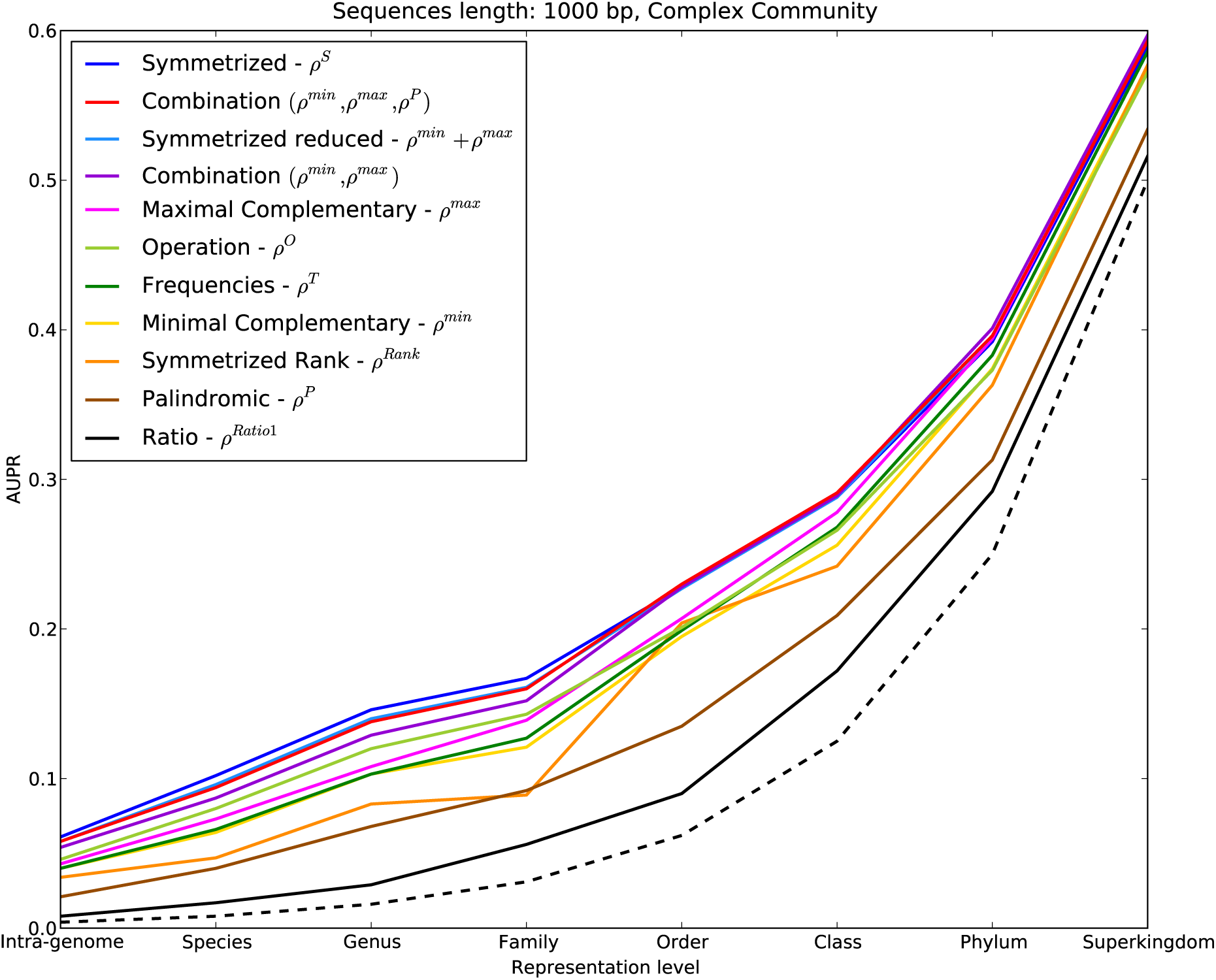
AUPRs obtained by the best signatures on complex community structure, for 1000 bp sequences. Dashed line is made by the AUPRs of a signature whose distance distributions are identical for each rank; the AUPRs of this signature do not depend on distribution shape. Signatures’ names in the legend are sorted with respect to the sum of their AUPRs for the different levels of taxonomic distance.

**Figure 3.**
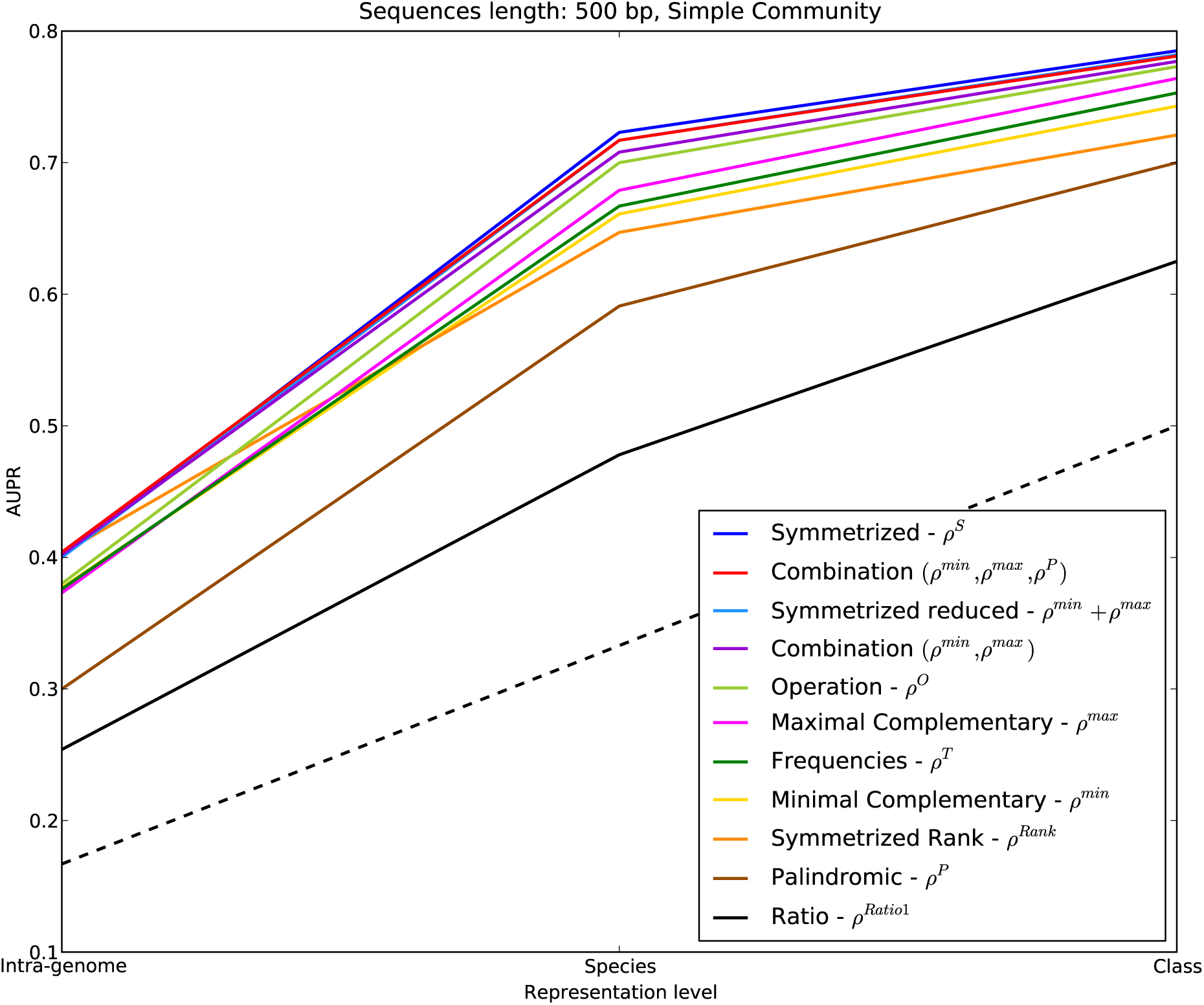
AUPRs obtained by the best signatures on simple-complexity community structure, for 500 bp sequences. Dashed line is made by the AUPRs of a signature whose distance distributions are identical for each rank; the AUPRs of this signature do not depend on distribution shape. Signatures’ names in the legend are sorted with respect to the sum of their AUPRs for the different levels of taxonomic distance.

**Figure 4.**
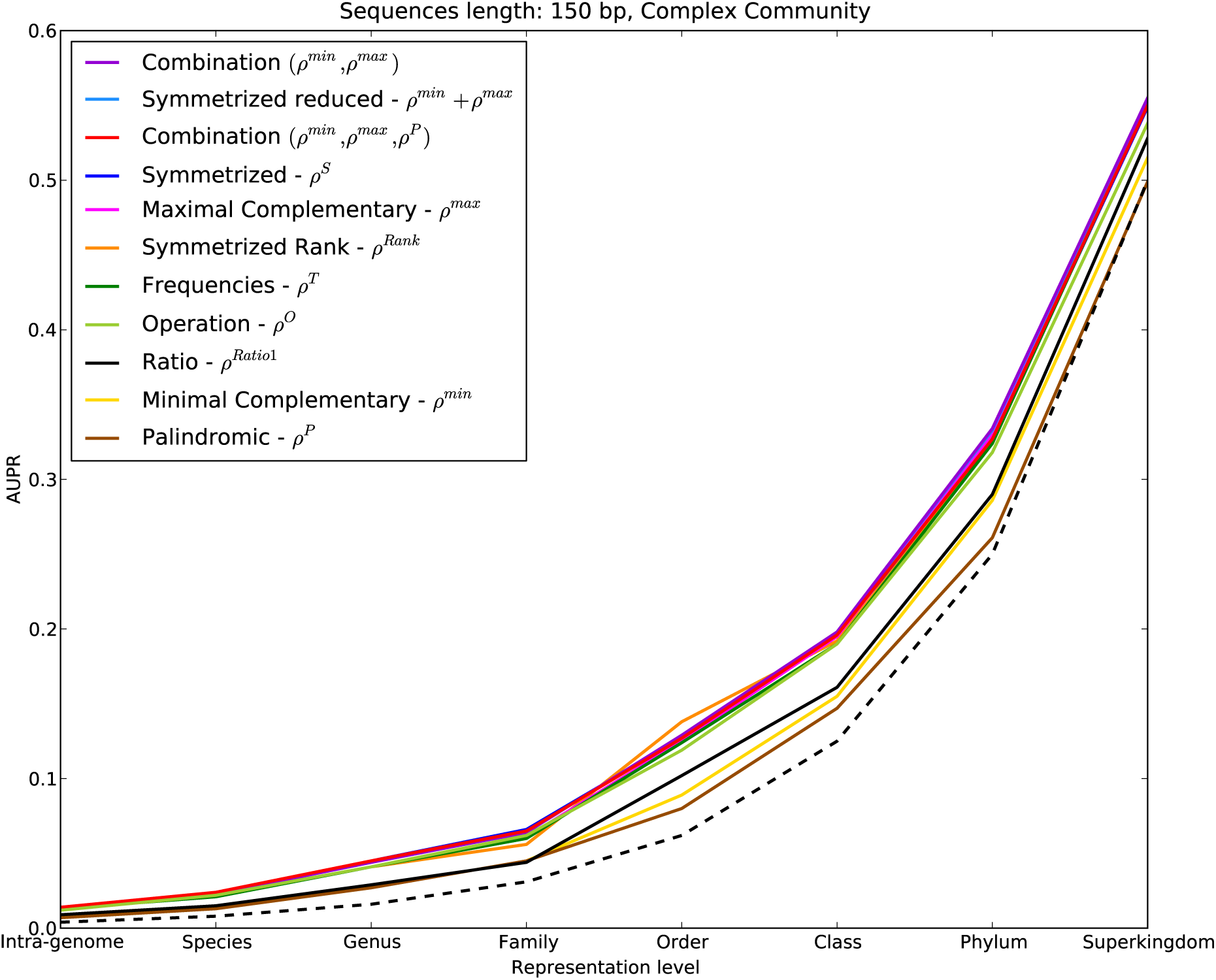
AUPRs obtained by the best signatures on complex community structure, for 150 bp sequences. Dashed line is made by the AUPRs of a signature whose distance distributions are identical for each rank; the AUPRs of this signature do not depend on distribution shape. Signatures’ names in the legend are sorted with respect to the sum of their AUPRs for the different levels of taxonomic distance.

The novel operation signature *ρ*^O^ was the signature with the lowest number of features among the ones superior or comparable to the standard signature *ρ*^T^ (*ρ*^T^ and *ρ*^O^ had 256 and 72 features, respectively). Theoretical analysis of *ρ*^O^ proved that it has the lowest intra-species dispersion among the signatures we study, but its inter-species discrimination power could be lower than the standard signature. Experiments indicate that its intra-species dispersion compensates its reduced inter-species discrimination power; this excellent intra-species dispersion allows operation signature to perform better than the standard signature. In agreement with theoretical analysis, experiments showed that operation signature gives the best performances when inter-species discrimination is easier. Adding frequencies of non-inverse k-mers probably leads to some information loss; however, the negative effect of this loss will be reduced when the microbial community is composed by few species. In this case, the likelihood of having organisms having different values for k-mer frequencies of reverse words *gi* and 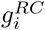 but the same value for the sum 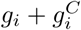 is reduced. Indeed, *ρ*^O^ excelled *ρ*^T^ for community structures of simple and medium complexity (Figure 3 and Supplementary Figure 4), while it had similar results for complex community structure (Figures 2 and 4). It is likely that *ρ*^O^ achieved good performances because it was inspired by a genomic rule [34], but we cannot exclude that similar results could be obtained by comparable reduction of *ρ*^S^(i.e. by summing frequencies of other sets of tetranucleotide frequencies that still lead to 72 features).

In accordance with our theoretical analysis, signatures capturing the deviation from GCSPR, like asymmetry signature *ρ*^A^, carried a phylogenetic signal; however, this signal was too weak to lead to appreciable performances. Figure 5 shows that the distance associated to signature *ρ*^A^ tends to assume slightly higher values for sequences coming from distantly related species; indeed, the signature distance distributions associated to higher levels of taxonomic distance are a bit shifted to higher values. Despite carrying a phylogenetic signal, signatures capturing the deviation from GCSPR were the worst in almost any experiment; ratio signature, which was the best deviation-capturing signature, had actually the worst Our theoretical analysis of ρ^A^ suggest performance among the signatures displayed in Figures 2 and 3. that this is due to high intra-species signature dispersion.

**Figure 5.**
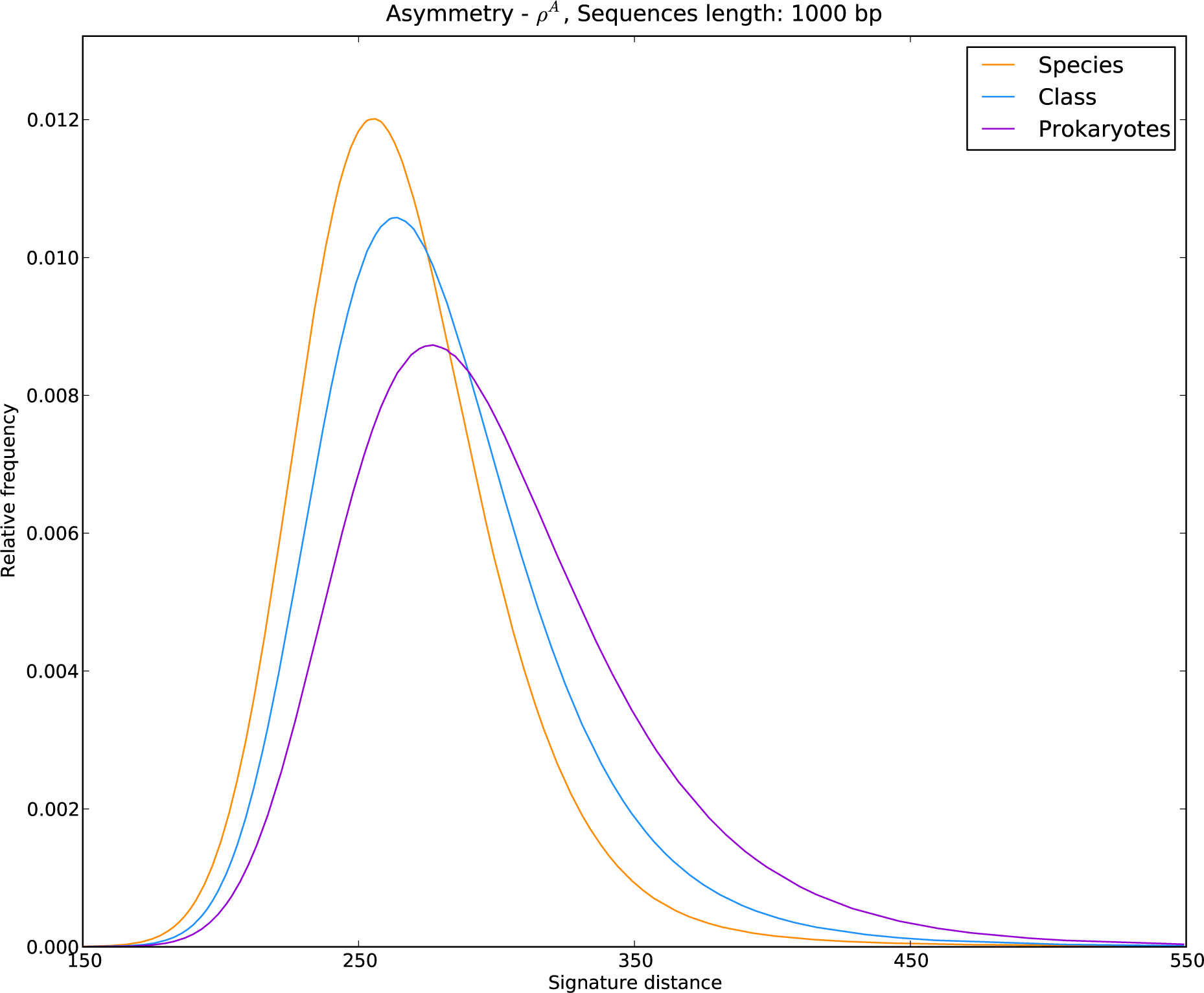
Normalized empirical distributions of Asymmetry Signature distances, for different levels of taxonomic diversity.

Signature performances increased with read length, as predicted by the theory. For instance, Figure 6 shows that longer sequences led to higher AUPRs for *ρ*^s^; the same happened for the other signatures. Theoretical analysis of symmetrized and standard signatures indicates that this is due to the reduction of intra-species dispersion, that tends to zero as read length *l* (and thus *n = l* – 4 + 1) increases – see Equations (8) and (6). The same observation holds for compositional, maximal complementary and minimal complementary signatures (11)-(12). This phenomenon is sensible also because the longer a sequences is, the more information it contains; hence, the compositional properties that characterize the source genomes are more recognizable. The trend was weaker for signatures designed to capture the deviation from GCSPR. In particular, their performances for complex community structure were not affected by sequence length (for *ρ*^Ratio1^, see Supplementary Figure 3). This could be due to the fact that their intra-species dispersion could be independent from read length; this is the case of the asymmetry signature (13) as our theoretical analysis showed.

**Figure 6.**
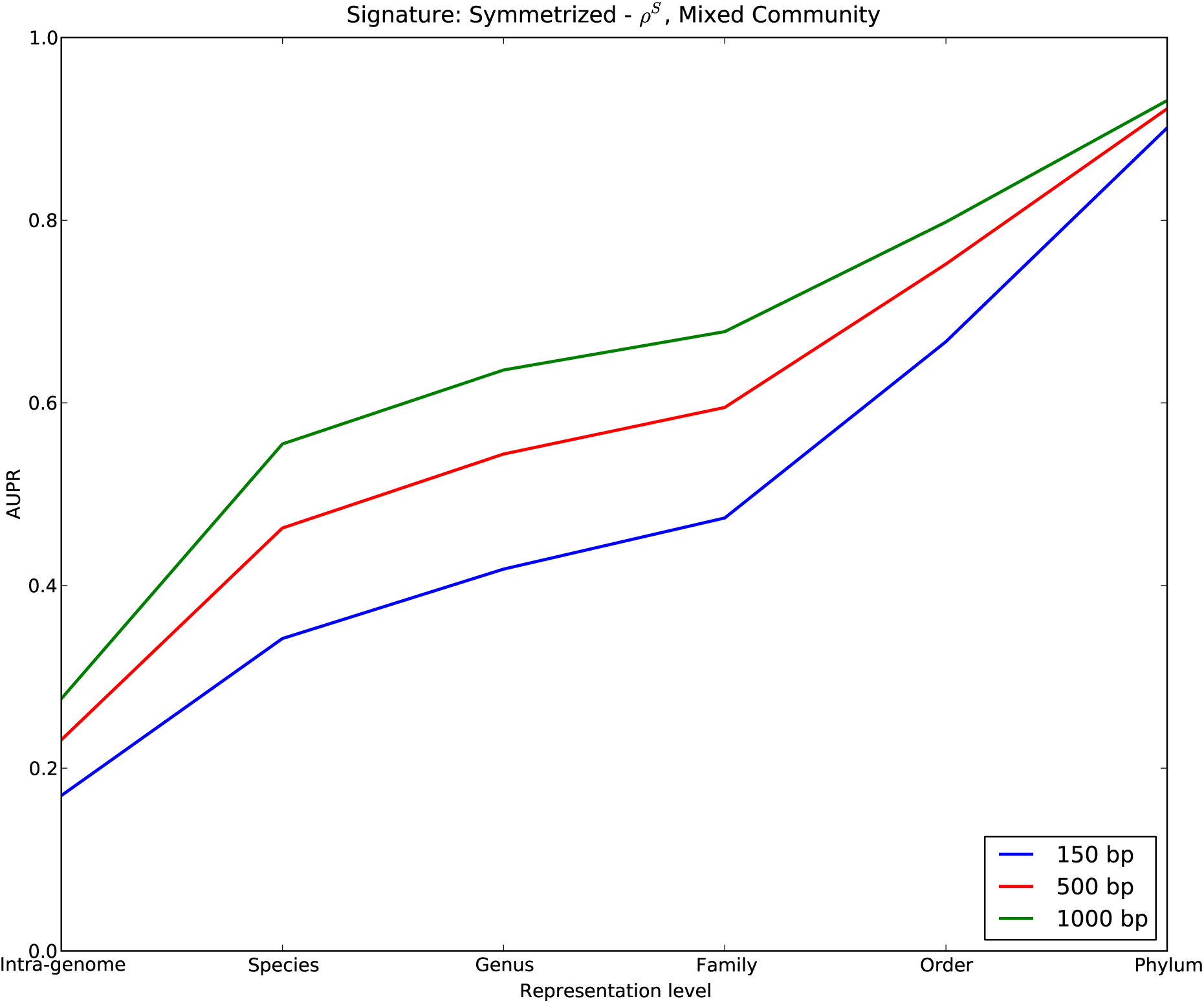
AUPRs obtained by Symmetrized Signature for all the sequence lengths. AUPRs were computed on medium-complexity community structure.

Reducing symmetrized and combination signatures to the features corresponding to the non-palindromic tetranucleotides worsened their performances, but they still outperformed the standard signature (Figure 7). Indeed, as mentioned before, signatures *ρ*^min^ + *ρ*^max^ and (*ρ*^max^, *ρ*^min^) outperformed the standard signature; moreover, they coincide with the features of *ρ*^S^ and (*ρ*^max^, *ρ*^min^, *ρ*^P^) corresponding to the non-palindromic tetranucleotides, respectively. Our theoretical analysis indicates that symmetrized and combination signatures outperform the standard signature thanks to their reduced intra-species dispersion; experimental results thus indicate that this factor is so strong that those two signatures outperform the standard signature even if they loose those 16 features of *ρ*^S^. Nevertheless, it might be possible that good performances could still be obtained by removing features corresponding to a different and perhaps larger set of *k*-mers.

**Figure 7.**
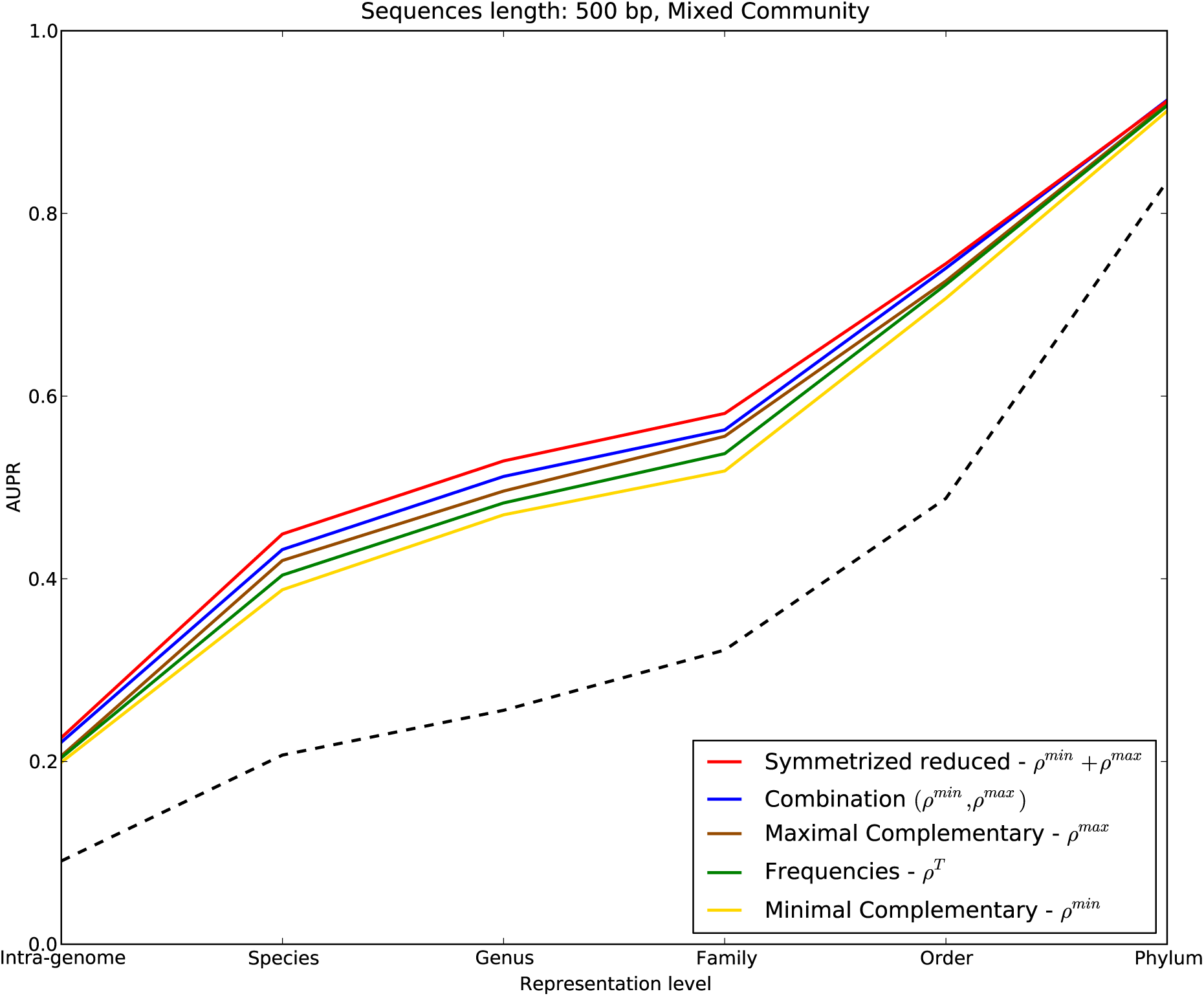
AUPRs obtained by a few asymmetry-related signatures on medium-complexity community structure, for 500 bp sequences. Dashed line is made by the AUPRs of a signature whose distance distributions are identical for each rank; the AUPRs of this signature do not depend on distribution shape. Signatures’ names in the legend are sorted with respect to the sum of their AUPRs for the different levels of taxonomic distance.

## 4 Conclusions

In this work, we conducted a theoretical analysis of new and existing genomic signatures for metagenomes; we analyzed their intra-species dispersion and their inter-species discrimination power. Furthermore, signatures’ performances were evaluated experimentally; the signatures were tested with respect to their capability to preserve the taxonomic relations of the source organisms of pairs of sequences. Signature distances were evaluated for sequences of 1,000 bp, 500 bp, and 150 bp randomly sampled from 1,284 prokaryotic genomes.

In metagenomic literature, symmetrized signature is erroneously believed to outperform the standard tetranucleotide signature simply because it is strand-independent. Our theoretical analysis proved that even the standard signature is actually not affected by the sampling strand; this is due to the GCSPR, which implies that frequencies in equally long reads of inverse k-mers follow the same probability distribution.

Moreover, our theoretical analysis showed that the success of symmetrized signature is due to its direct exploitation of GCSPR. By summing frequencies of inverse k-mers, the inter-species discrimination power is unmodified: there is no information loss because the frequencies of the combined k-mers follow the same probability distribution. Moreover, this summation reduces the intra-species dispersion, thus leading to performance improvement. Experimental results confirmed that symmetrized signature had the best performance among all the tested signatures; the dimension of its feature space is about half of the one of the standard signature, making the data also more tractable.

On the contrary to what is believed, summing frequencies of inverse k-mers is not the only effective way to produce an effective strand-independent signature from the standard signature. The novel combination signature achieves strand-independence through a biologically-sensible reordering of standard signature’s features. As predicted by the theory, experimental performances of combination signature were a bit below symmetrized signature but still higher than standard signature. Combination signature thus seems the ideal choice if oligonucleotide frequencies must be kept separated; in general, it could be used by alignment-free sequence comparison methods when source genomes do not respect GCSPR or distinguishing inverse oligonucleotides matters. In particular, it can be advantageous for tasks on metagenomic data that are not strictly related to taxonomy, like the identification of cis-regulatory modules [21]. It can still be helpful for taxonomic analysis of metagenomes when they contain sequences sampled from genomes not respecting GCSPR, like viral genomes. Experimental results also indicate that some features of symmetrized and combination signatures can be removed without too much decrease of performance.

The performances of the novel operation signature indicates that other genomic symmetries than GCSPR can be successfully exploited for designing low-dimensional signatures. Indeed, despite having lower performance than the symmetrized signature, it has about half of its features and was superior to the standard signature for communities with not-very-complex taxonomic structures (and at least comparable for very complex ones). Theoretical analysis proved that it has the lowest intra-species dispersion among the studied signatures; however, its inter-species discrimination power could be lower of the one of the standard signature. Therefore, operation signature can be particularly beneficial for the analysis of large metagenomes sampled from microbial communities whose taxonomic structure is not extremely complex; indeed, dealing with less features reduces the computational cost of data analysis [36].

Deviation from GCSPR seems related to the taxonomic classification of the species, but not strongly: signatures that are exclusively based on these asymmetries are not good enough to achieve the best performances. Theoretical analysis of asymmetry signature suggests that performances are low due to a high intra-species signature dispersion.

As predicted by the theoretical analysis and confirmed by the experiments, signature performances increase with sequence length. It is likely that no signature can be effective on very short sequences, due to the little information contained. Some works [16,25,26] dealt successfully with this issue by adopting the following approach: short sequences were grouped, and then the signature was computed on this set as if it were one long sequence. However, this procedure might be risky for metagenomes with high diversity or low species coverage [52]. An alternative method is to compute the signatures on reads assembled into contigs [18] but unfortunately the assembly process of a metagenome is computationally very intensive.

Our approach provides a unified framework to compare the performances of signatures by means of binomially and normally distributed random variables. This framework can be used to guide the development of novel signatures.

## Acknowledgments

FG participated in conceiving the study, performing the data analyses, and interpreting the results; he developed the theoretical statistical analysis, generated the data, and wrote the manuscript. DM participated in conceiving the study, analyzing the data, interpreting the results, and supported the writing of the manuscript. MJ participated in conceiving the study and in interpreting the results. EM participated in conceiving and coordinating the study, analyzing the data, and helped writing the manuscript. All authors read and approved the final manuscript.

## Footnotes

1 In those works, distance between sequences was measured through Spearman footrule distance, that is equivalent to compare the values assumed by ρ^Rank^ via L_1_.

2 Available at ftp://ftp.ncbi.nih.gov/genomes/Bacteria/

3 Available at http://cs.ru.nl/~gori/download/Table_S1_list_genomes.txt

4 Available at ftp://ftp.ncbi.nih.gov/pub/taxonomy/

